# Steric Interactions at Gln154 in ZEITLUPE Induce Reorganization of the LOV Domain Dimer Interface

**DOI:** 10.1101/2020.10.05.326595

**Authors:** Ashutosh Pudasaini, Robert Green, Young Hun Song, Abby Blumenfeld, Nischal Karki, Takato Imaizumi, Brian D. Zoltowski

## Abstract

Plants measure light, quality, intensity, and duration to coordinate growth and development with daily and seasonal changes in environmental conditions, however, the molecular details linking photochemistry to signal transduction remain incomplete. Two closely related Light, Oxygen, or Voltage (LOV) domain containing photoreceptor proteins ZEITLUPE (ZTL) and FLAVIN-BINDING, KELCH REPEAT, F-BOX 1 (FKF1) divergently regulate the protein stability of circadian clock and photoperiodic flowering components to mediate daily and seasonal development. Using structural approaches, we identified that mutations at the Gly46 position led to global rearrangements of the ZTL dimer interface. Specifically, introduction of G46S and G46A variants that mimic equivalent residues found in FKF1 induce a 180° rotation about the dimer interface that is coupled to ordering of N- and C-terminal signaling elements. These conformational changes hinge upon rotation of a C-terminal Gln residue analogous to that present in light-state structures of ZTL. The results presented herein, confirm a divergent signaling mechanism within ZTL that deviates from other members of the LOV superfamily and suggests that mechanisms of signal transduction in LOV proteins may be fluid across the LOV protein family.

## Introduction

Light, Oxygen, or Voltage (LOV) domain containing proteins are abundant in nearly all kingdoms of life, where they function as sensory proteins to regulate a diverse array of adaptive responses^*1, 2*^ Structurally, LOV proteins consist of a central 5-stranded β-sheet flanked on one side by four helical elements that cradle a photoactive flavin-based cofactor (FAD, FMN and riboflavin)^*1, 3*^ (Fig. 1). Upon photoexcitation a covalent bond is formed between a conserved cysteine residue and the C4a position of the flavin cofactor (C4a adduct)^*4-7*^. The C4a adduct is reversible in the presence of UV-light, or will spontaneously decay in the dark with variable kinetics ranging from seconds to days^*4, 6, 8*^. The result is a tunable photochemical sensor that can modulate activity over a wide range of environmental light intensities^*9, 10*^.

**Fig. 1.**
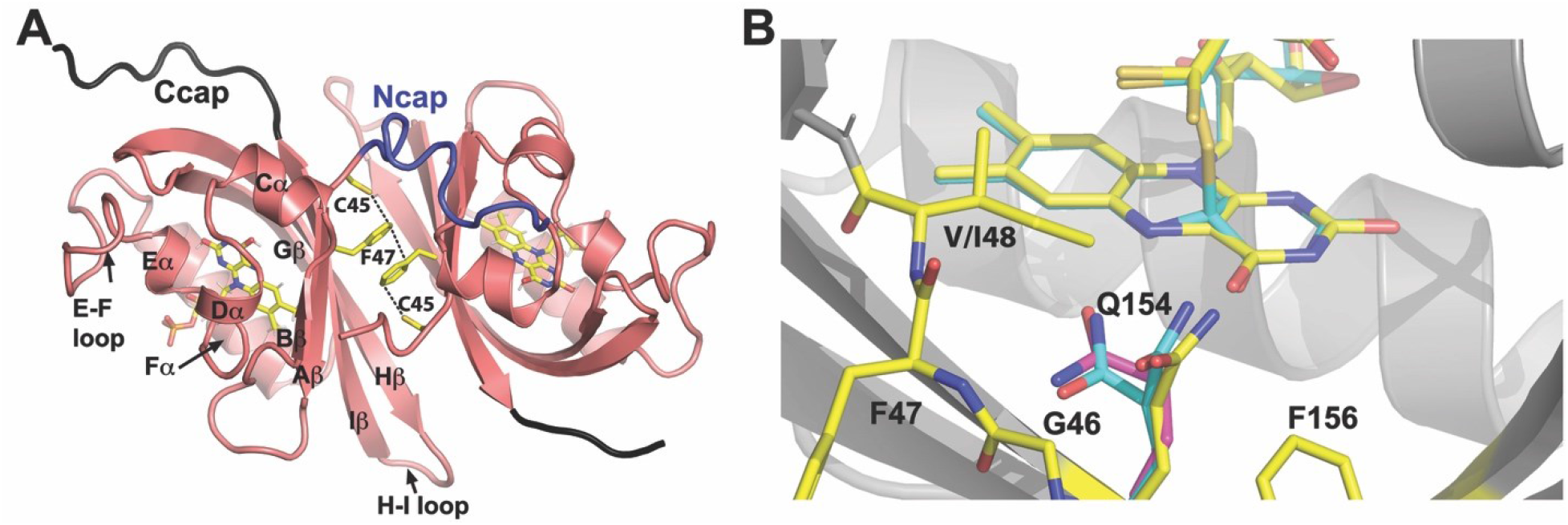
Structure and signaling mechanisms of ZTL. A) Previous ZTL crystal structures consist of an antiparallel dimer mediated by extensive contacts along the β-sheet surface. A network sulfur-π and π-π interactions between Cys45 and Phe47 within the Ncap region stabilize the dimer interface. B) Crystal structures of ZTL demonstrated an unusual exposed orientation of Gln154 in the dark (yellow), compared to a buried orientation in all other LOV structures (magenta). Photoactivation in a V48I:G80R variant leads to formation of a C4a adduct (cyan) and sp^2^-sp^3^ hybridization of C4a and protonation of the N5 position. Light-state structures of a V48I:G80R variant indicate that Gln154 may rotate to a buried position following adduct formation.

Structural and computational studies have linked C4a adduct formation to global conformational changes in protein structure. Existing studies focus on two elements of photoactivation, a change in the hybridization state of the C4a position of the flavin isoalloxazine ring following adduct formation, and protonation of the adjacent flavin N5 position^*11-14*^. Light- and dark-state structures of LOV proteins confirm that N5 protonation causes a flip in the orientation of a conserved Gln residue that alters a hydrogen-bonding network connected to the β -sheet surface. Alteration in H-bonding interactions dictates either reorganization of the β -sheet surface, or alterations in the conformation of N- and C-terminal extensions to the LOV core^*12, 15*^ (Fig. 1). These downstream conformational changes then initiate signal transduction through rearrangement of protein-protein interaction networks or relay photochemical signals to N- or C-terminal signal transduction domains to modulate enzymatic activity in response to changes in blue-light intensity^*1, 16-19*^. In the latter case, the types of signal transduction domains are diverse, leading to LOV domains being viewed as a plug-n-play module that can impart photoactivity to enzymatic function^*1, 2, 19*^.

Recent research has indicated that N5 protonation is both necessary and sufficient for photochemical activation. First, studies of LOV variants, which cannot form the C4a adduct, but retain light-driven photoreduction, are photochemically active and competent for signal transduction^*13*^. Second, Gln→Leu substitutions that would retain an ability to sense alterations in steric interactions following a hybridization change abrogate signal transduction in most characterized LOV proteins^*20, 21*^. However, the universality of a Gln-flip mechanism was recently called into question by structures of the plant circadian clock photoreceptor ZEITLUPE (ZTL)^*9*^

In plants, the ZTL family of photoreceptors consists of ZTL, FLAVIN-BINDING, KELCH REPEAT, F-BOX 1 (FKF1), and LOV KELCH PROTEIN 2 (LKP2). All three are modular LOV proteins, consisting of an N-terminal LOV domain that imparts blue-light regulation of C-terminal F-box and Kelch repeat domains^*22, 23*^. Absorption of blue-light regulates two primary activities: 1) Light-driven formation of a ZTL-GIGANTEA (GI) complex, which results in mutual stabilization and 2) Light-driven regulation of E3-ligase activity to allow degradation. Notably, despite conservation of GI-complex formation, ZTL and FKF1 differ in the timing of degradation of protein targets, whereby light inhibits ZTL mediated degradation of circadian clock components, which is enhanced following nightfall^*24-26*^. In contrast light activates the E3 ligase activity of FKF1 allowing enhanced targeted degradation during the day.

Structural studies of ZTL identified alteration in the canonical mechanism of LOV signal transduction. Namely, structures indicated that the active site Gln residue that is essential for function adopts a heterogeneous conformation under dark-state conditions. Light-state crystal structures revealed that adduct formation led to rotation of Gln154 from an exposed conformation to a buried conformation consistent with all other LOV domain structures (Fig. 1B)^*9*^. These structural studies revealed that heterogeneity in Gln154 conformation was facilitated by the presence of Gly46, which evolutionarily differentiated ZTL from FKF1 and other plant LOV proteins. Unfortunately, these structural studies were based on a ZTL variant containing a V48I substitution that disrupts complete rotation of Gln154, thereby blocking full population of the light-state and impeding identification of any global alterations in protein structure. In these regards, the light-state structures are unable to capture the global conformational response attenuating the E3-ligase activity of the effector domains, and also unable to identify how signaling mechanisms may differ in ZTL and FKF1 to allow differential function depending on time of day. Herein we employ structural approaches to identify an allosteric mechanism regulating reorganization of the ZTL dimer that is dependent on the residue identity at Gly46. These results confirm an alternative mechanism of signal transduction in ZTL and indicates that LOV allostery may be more fluid across the LOV superfamily.

## Materials and Methods

### Cloning and Purification

ZTL G46S:G80R and G46A:G80R constructs composed of residues 29-165 were cloned, expressed and purified with a GST or 6xHIS fusion tag and purified as reported previously^*9*^ All proteins were purified in 50 mM Tris pH 7.4, 100 mM NaCl, and 10% glycerol (Buffer A). Cells were lysed via sonication at 4°C. After sonication the cell debris was separated from the lysate by centrifugation at 18,000 RPM at 4°C for 60 min. Depending on the affinity tag, the proteins were purified using either Ni-NTA or GST affinity columns. The proteins were incubated overnight at 4°C with 6His-TEV protease to cleave the affinity tags. Cleaved affinity tags and the TEV protease were separated on a Ni-NTA column followed by a final round of purification using Fast Protein Liquid Chromatography (FPLC) on a Hiload Superdex 16/60 gel filtration column equilibrated with Buffer A.

### Structural analysis

ZTL crystals were initially obtained from Hampton Screens (HR2-110 and HR2-112) via hanging drop method using 1.5 μL of crystallization solution with 1.5 μL of ZTL variant at a concentration range of 5-10 mg/mL. Optimum crystallization conditions were determined for G80R:G46S (0.05 M Tris pH 8.5, 2.0 M Ammonium Sulfate), and G80R:G46A (0.1 M MES pH 6.0, 1.6 M Ammonium Sulfate, 0.01 M Cobalt Chloride Hexahydrate). Protein for crystallographic studies was purified in Buffer A. Crystal trays were set in a dark room illuminated with a red light.

Diffraction data was acquired at the F1 beamline at the Cornell High-Energy Synchrotron Source (CHESS). All data was collected at 100 K. The following cryoprotectants were added: G80R:G46S (25% ethylene glycol v/v %), and G80R:G46A (25% ethylene glycol v/v %). Data reduction and scaling was done in HKL2000^*27*^. ZTL variants were solved using molecular replacement in PHASER^*28*^ with WT ZTL as the search model. Rebuild cycles were performed in COOT^*29*^ and the data was refined with PHENIX^*30*^. Coordinates for G46S:G80R (PDB: 6WLP) and G46A:G80R (PDB: 6WLE) structures can be found in the protein data bank.

### SEC-SAXS Sample Preparation and Analysis

All Size-exclusion chromatography-Small angle X-ray Scattering (SEC-SAXS) samples were purified as outlined in cloning and purification with the exception that the final purification buffer contained 50 mM HEPES pH 8.0, 100 mM NaCl, and 2 mM TCEP, prior to concentration to 20 mg/ml. SEC-SAXS data were collected at Cornell high energy synchrotron source (CHESS) on beamline G1. Prior to data collection, all samples were centrifuged at 14,000 RPM for 20 minutes. SEC was conducted at 4°C using a Superdex 200 5/150 column on an AKTA Pure System (GE Healthcare Life Sciences, Marlborough, MA) with a flow rate of 0.15 mL/min and 0.5 s exposures. The column was equilibrated with 50 mM HEPES pH 8.0, 100 mM NaCl, and 2 mM TCEP. SAXS data were collected at 9.9099 keV (1.2511 Å) at 7.19×1011 photons/s. The X-ray beam was collimated to 250 × 250 μm^2^ diameter and centered on a capillary sample cell with 1.5 mm path length and 25 μm thick quartz glass walls (Charles Supper Company, Natik, MA). The sample cell and full X-ray flight path, including beamstop, were kept in vacuo (< 1×10-3 torr) to eliminate air scatter. Temperature was maintained at 4°C. Images were collected on a dual Pilatus 100K-S detector system (Dectris, Baden, Switzerland).

Data processing, image integration, normalization, subtraction, and merging were conducted in BioXTAS RAW 2.0.1^*31*^. Data was further modeled using software available in the ATSAS software package^*32*^. Pairwise distribution and Guinier analysis was conducted with GNOM. The *ab initio* molecular envelopes were generated using DAMMIF with P1 symmetry. 35 *ab initio* models were constructed, aligned and averaged using DAMAVER, and refined with DAMMIN. The mean normalized spatial discrepancy (NSD) across all models was determined to be 1.118+/-0.115. One model was found to have an NSD in excess of the mean NSD +/- 2*SDV and thus was excluded from the final averaged envelope. The final reported envelope demonstrates a chi^2^=1.191. SAXS data has been deposited in SASBDB (SASDJXS).

### Protein-Protein Interactions Assay

To overexpress 3xFLAG-6XHis (3F6H) tag fused ZTL variant proteins, the full-length *ZTL, ZTL G46A, ZTL G46S*, and *ZTL Q154L* cDNAs (originally cloned in pENTR/D-TOPO) were introduced into pB7-HFN binary vector^*33*^. To analyze protein-protein interactions, the *35S:3F6H-ZTL, 35S:3F6H-ZTL G46A, 35S:3F6H-ZTL G46S, 35S:3F6H-ZTL Q154L* and *35S:HA-GI* constructs were infiltrated into 3-week-old *N. benthamiana* leaves as described^*9*^.

Co-immunoprecipitation (Co-IP) assays were performed as described to test protein-protein interactions between ZTL variants and GI^*34*^. Briefly, the leaf tissues were harvested and ground in liquid nitrogen. Proteins were extracted from 0.5 ml volume of the ground tissues using Co-IP buffer [50 mM Na-phosphate pH7.4, 150 mM NaCl, 10% glycerol, 5 mM EDTA, 1 mM DTT, 0.1% Triton X-100, 50 μM MG-132, 2 mM NaVO4, 2 mM NaF, and protease inhibitor tablets-EDTA free (Pierce)]. The 3F6H-fused ZTL variants were precipitated (at 4 °C for 10 minutes under dim light) with anti-FLAG antibody (Sigma), which was bound to Protein G-coupled magnetic beads (Dynabeads Protein G, Invitrogen). Precipitated proteins were resolved in 8% SDS-PAGE gels, and the presence of ZTL variants and HA-GI was detected by western blot using anti-FLAG (Sigma) and anti-HA (3F10, Roach) antibodies, respectively.

## Results

Previous studies of the ZTL family identified the LOV domains as obligate dimers formed by extensive contacts along the β-sheet surface^*9*^. In FKF1, solution scattering experiments indicated that light induces reorganization of an anti-parallel FKF1 dimer interface in a manner dependent on the E-F loop^*35*^, however structural details regarding the nature of the conformational response is not known. In ZTL, dark- and light-state structures identified two similar anti-parallel dimers mediated by contacts across the central β-sheet, that differ only by a 2.0 Å translation along the dimer interface^*9*^. The role of either of these two dimers in signal transduction is currently unknown. Further, how ZTL and FKF1 may differ at the level of signal transduction is poorly understood.

Previous studies indicated that ZTL has an unusual mechanism of signal transduction that may stem from the evolutionary selection of a Gly residue at position 46. Phylogenetic analysis indicated the residue identity at this position differentiates ZTL (Gly) from other plant LOV proteins, such as FKF1 (Ala or Ser) and phototropins (Asn). Thus, we sought to obtain the crystal structures G46S and G46A variants of the isolated ZTL LOV domain to examine potential structural and functional differences. In both cases, proteins had to be studied in the background of a G80R variant that enhances stability, but does not affect protein structure. Similar to WT-ZTL, G46A and G46S variants are constitutively dimeric in both the dark- and light-states, suggesting no gross perturbation in structure^*9*^.

Examination of the dimer interface in the 3.0 Å crystal structures of G46A and G46S reveal distinct structural differences that result from reorientation of Gln154 (Fig. 2). As detailed below, reorientation of Gln154 impacts local structure at an N-terminal CGF (Cys45-Gly46-Phe47) and a C-terminal QFF (Gln154-Phe155-Phe156) motif that are coupled to global reorientation of the dimer interface. Below, we focus on specific structural differences, beginning at the level of global crystal packing.

**Fig. 2:**
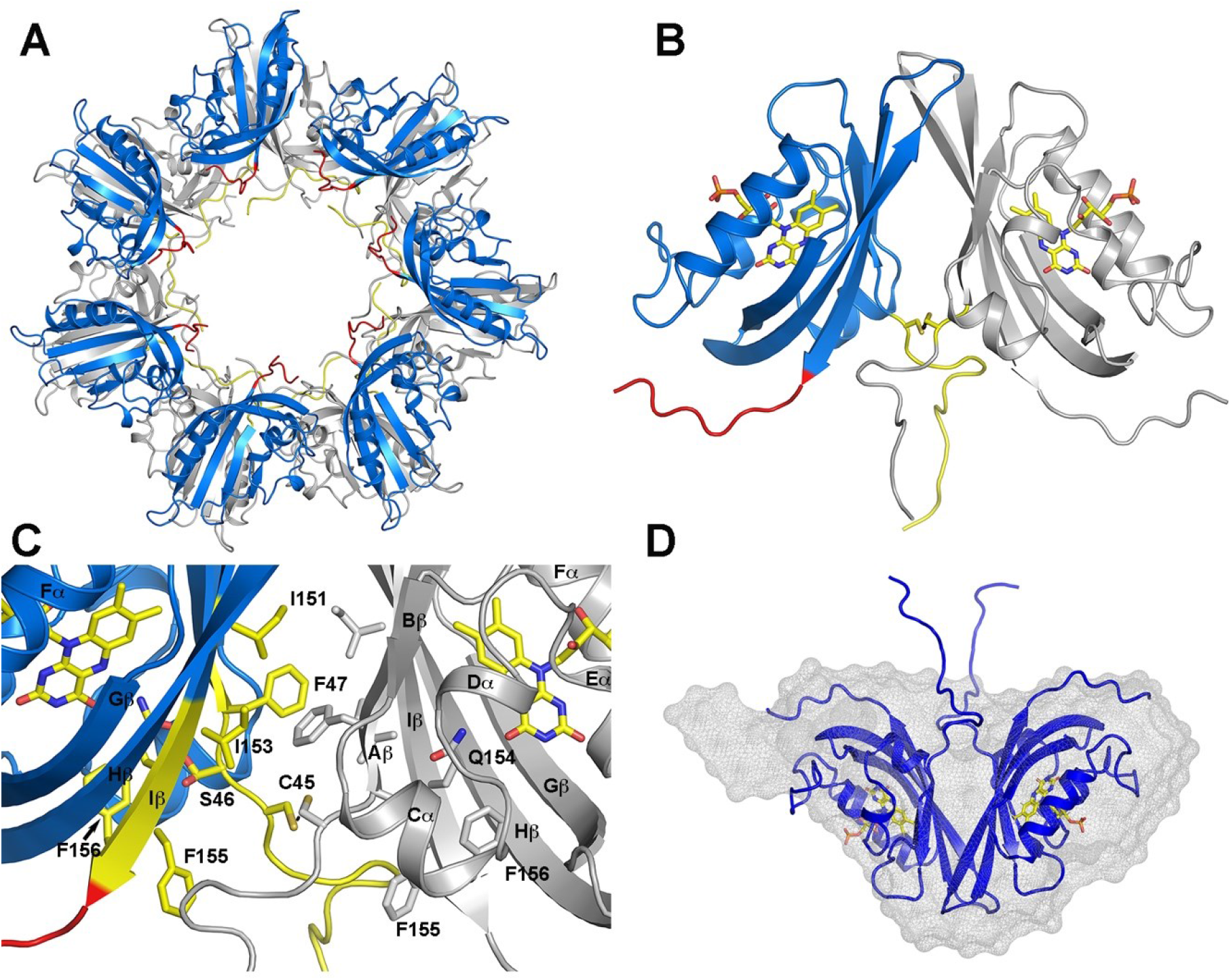
G46A/S induces global reorganization of the dimer Interface. A) G46A/S:G80R crystalize in spacegroup I213 as a 7-fold symmetric ring (blue). The two-fold screw axis generates a second copy of the 7-fold ring, linked by disulfide linked dimers in all 7-copies B). The crosslinked dimer (B) is parallel in orientation and leads to ordering of the N- and Ccaps. C) The parallel dimer is linked through Cys45. Hydrophobic contacts along the β-sheet are similar to WT ZTL, involving Phe47, Ile151 and Ile153, but absent the distinguishing sulfur-π and π-π interactions. D) SAXS of G46S:G80R variants generates an envelope (gray mesh) consistent with the crystallographic dimer.

Although the core LOV structures of G46A and G46S are highly similar to WT-ZTL, reorganization of N-terminal and C-terminal elements leads to crystallization in an alternative space group (I213 vs. P3121:WT) and grossly alters crystal packing interactions resulting in dimeric, tetrameric, and heptameric assemblies depending on which molecules are selected in the asymmetric group (Fig. 2 and Fig. S1). Both G46A and G46S crystalize in a 7-fold symmetric ring involving seven copies of the isolated LOV domain (Fig. 2A). Ncap and Ccap elements are well defined in both structures and orient into a central cavity. In addition, the two-fold screw axis generates a second copy of the symmetric ring that forms dimeric contacts between individual LOV domains. These dimeric contacts involved the central β-sheet similar to WT-ZTL, however instead of an antiparallel arrangement observed in WT proteins, Gly46 variants adopt a parallel dimer (Fig. 2B,C). Notably, SEC-SAXS confirms G46S:G80R exists as a dimer in solution (MW=30.8 kDa; Rg=22.17 Å). Moreover, *ab initio* reconstructions of the molecular envelope are consistent with the crystallographic dimer (Fig. 2D and supplementary Fig. S2). Thus, introduction of Gly46 variants leads to a 180 degree rotation about the β-sheet interface that is coupled to ordering of N-terminal and C-terminal elements.

Although the primary dimeric interface in G46A and G46S still involves the β-scaffold, contacts mediated by the PAS β-sheet deviate from WT-ZTL (Fig. 2B, C). Initial inspection of the β-scaffold dimer reveals interactions similar to WT ZTL. The β-sheet directed dimer is stabilized by extensive hydrophobic contacts involving Phe47, Ile151 and Ile153, consistent with WT proteins (Fig. 2C). However, the parallel orientation abolishes the sulfur-π and π-π network present in WT-ZTL. Instead, the parallel orientation is facilitated by disulfide bond formation between Cys45 residues (Fig. 2B,C). Importantly, treatment of G46A/S with TCEP during SEC-SAXS has no effect on dimer formation by SEC (Fig. S2). Thus, these disulfides are not required for dimer formation; rather, they likely result from the close interaction of Cys45 that can trap the parallel dimer.

Analysis of the intersection of Gln154 with the CGF and QFF motifs identifies conformational changes facilitating rearrangement of the dimer interface. In both G46A and G46S introduction of a side chain leads to a steric clash with exposed conformations of Gln154. These steric contacts push the Ncap away from Iβ allowing formation of the Cys45 disulfide bond and disruption of the sulfur-π interaction network involving Cys45 and Phe47 within the CGF motif (Fig. 3). Disruption of the sulfur-π interaction network stabilizing the anti-parallel dimer in leads to a 180 degree rotation about the WT dimer interface.

**Figure 3.**
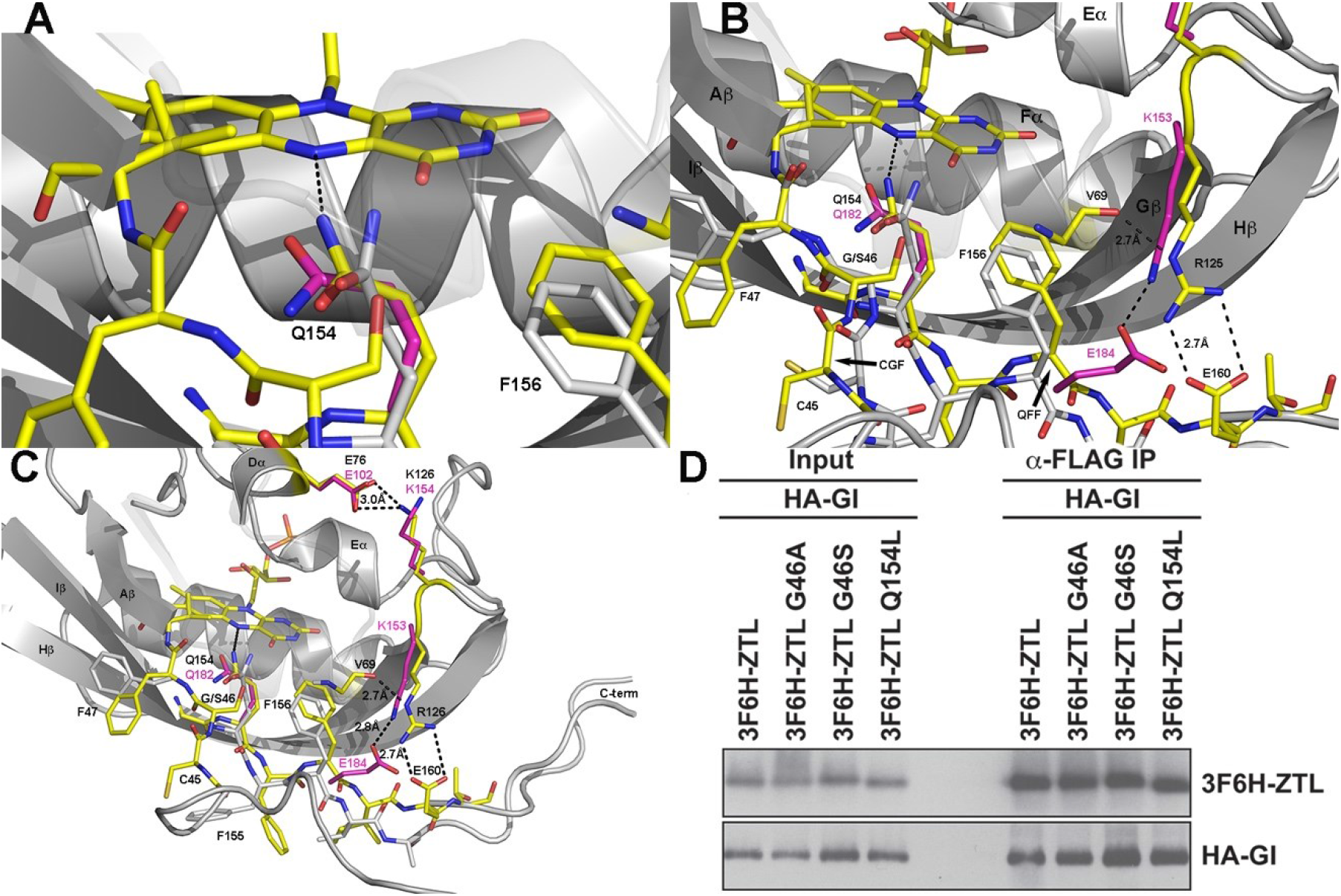
Gln154 reorientation dictates N- and C-termini conformation. A) Introduction of G46S or G46A variants (yellow) leads to rotation of Gln154 towards the buried conformation (magenta) compared to the exposed conformation (grey). Rotation of Gln154 is coupled to movement of Phe156 towards the FMN active site. B, C) Comparative structures of G46S (aqua), WT ZTL (yellow) and VVD (magenta). The altered interactions in G46S are linked to a buried conformation of Q154 that propagates to the Ncap, Ccap, and helical interfaces. These changes are coupled to formation of a salt bridge between Glu158 and Arg125 that are conserved in diverse LOV proteins (VVD shown in magenta). These further propagate from the G-H loop (Arg125, Lys126) to the helical interface through a conserved salt bridge to Glu158. D) Confirmation of protein-protein interactions between ZTL variants and GI *in planta*. ZTL variants were immunoprecipitated by anti-FLAG antibody and HA-tagged GI protein was detected by anti-HA antibody. The similar results were repeated two times independently, and the representative image is shown.

In concert with movements at the N-terminal CGF motif. Steric interactions between position 46 and the active site Gln154 results in Gln154 rotation toward the buried conformation observed in light-state ZTL (Fig. 3A-C). Rotation of Gln154 induces movement of Phe155 and Phe156 to partially occupy the region voided by Gln154. These movements draw C-terminal elements closer to the LOV core, partially extending β-sheet-like contacts at the C-terminal end of Iβ. Ordering of the C-terminus results in formation of a salt bridge between Arg125 within the G-H loop and Glu158 within the Ccap (Fig. 3B,C). Further, the G-H loop makes extensive contacts with the Cα helix. These contacts primarily involve two adjacent, positively charged residues Arg125/Lys126. In addition, to forming a salt bridge with Glu158, Arg125 forms a H-bond to the carbonyl moiety of Val69 in the Cα helix. The neighboring Lys126 forms an additional salt bridge to Glu76 at the C-terminal end of the D-helix. In this manner, Glu158 anchors Iβ to the LOV core allowing the Cα helix to form a cap directly above the QFF and CGF motifs (Fig. 3C). Thus, rotation of Q154 propagates throughout the Ncap and Ccap elements to reorganize the LOV-dimer interface and stabilize interactions between the Ccap and LOV core through new salt-bridge interactions. The net result is a more stably folded LOV domain with clear density for both the N/Ccap.

The results outlined above, suggest a divergent signal transduction mechanism in ZTL, whereby the primary signaling element involves both steric interactions between Gln154 and the flavin O4 position, as well as alteration in H-bonding networks initiated from N5 protonation. If steric interactions stemming from a change in hybridization of the C4a position are a driving factor for light-driven changes in ZTL structure, then Q154L variants may be competent for signal transduction, in contrast to other characterized LOV proteins. To evaluate the necessity of Q154 in ZTL signaling we examined the effect of Q154L, G46S and G46A variants on light-activated complex formation with GI in tobacco. Consistent with our hypothesis, all three variants robustly form the ZTL-GI complex following exposure to light (Fig. 3D). Thereby, confirming Q154L variants are competent for light-state complex formation with GI and that steric interactions alone are sufficient for light-induced ZTL-GI complex formation.

## Discussion

Previous structural studies of the ZTL family of LOV photoreceptors indicated that the signal transduction mechanism of ZTL diverges from other LOV proteins, and that photoactivation leads to rearrangement of a dimer interface^*9, 23, 35*^. However, the precise nature of the global reorganization remained elusive due to either the lack of atomic resolution crystal structures (FKF1) or the presence of mutations that block global reorganization of the protein (ZTL). Herein, we have identified a protein variant that traps a light-state-like orientation of the active site Gln residue in crystal structures. The resulting structures allow a snapshot of global reorganization following rotation of Gln154 from the exposed (dark) to buried (light-like) conformation. The new crystal structures shed light on ZTL function and signal transduction within the LOV superfamily.

Comparison of the ZTL crystal structures identifies a conserved topology of signal transduction in LOV proteins. First, photoactivation results in two primary signal-inducing events, protonation of the N5 position and a sp^2^ to sp^3^ change in hybridization at the C4a position^*6, 12, 36, 37*^. Second, studies of the LOV protein VVD and a engineered LOV histidine kinase, YF1, confirmed that N5 protonation was both necessary and sufficient for photoactivation^*13*^. Here, we show that ZTL adopts an alternative signaling mechanism, whereby an H-bonding residue at the Gln154 position is not necessary, and light-state functionality is retained in a Q154L variant. We propose that in ZTL, steric interactions due to C4a hybridization contribute to ordering Gln154 and thereby stabilizing the buried conformation. Ordering of Gln154 in a buried conformation leads to movement of F156 to fill a position vacated by Gln154 resulting in ordering of the C-termini and induction of a stabilizing salt-bridge. In concert with C-terminal conformational changes, the Aβ-strand moves away from the LOV core destabilizing the anti-parallel dimer interface and leading to a 180° rotation about the central β-scaffold.

### Implications for LOV Signal Transduction

Comparisons of the ZTL structures to recent simulations of VVD indicate convergence of a previously uncharacterized motif in LOV allostery. In VVD, light-activation also leads to increased stability of a C-terminal salt-bridge^*38*^. Further, residues capable of forming the Arg125 -Glu158 salt bridge are retained in existing LOV structures, suggesting the signaling motif may be conserved across the LOV superfamily. In these regards, the initial driving force of an allosteric response (hybridization or H-bonds) may diverge in LOV proteins, however they conserve signaling elements that enable order-disorder transitions at the N- and C-termini, that hinge upon residue selection at a key allosteric signaling site (Gly46).

### Effects on ZTL Signal Transduction

ZTL crystal structures confirm that the alteration in the ZTL signaling mechanism stems from the unusual evolutionary selection of Gly46. The equivalent residue is a Ala/Ser in fungal LOV and in plant FKF1 proteins, or an Asn residue in plant phototropins^*9*^. In these cases, the residue occupying that position is involved in alterations in H-bonding interactions following the Gln-flip mechanism, enabling adduct formation to signal to the N- and C-termini of LOV proteins^*9, 12, 14, 36*^. In ZTL, the Gly residue is unable to respond to H-bond changes, rather, the lack of a side chain permits alternative conformations of Gln154, enabling a hybridization-sensing mechanism to proceed. Introduction of an Ala/Ser residue in ZTL introduces steric constrains on Gln154 invokes a light-state like rearrangement of the dimer interface by promoting concerted movements of Gln154, the N-terminal CGF motif, and the C-terminal QFF motif. These observations let us to postulate on signaling differences between ZTL and FKF1 proteins in plant photobiology.

Biological activity of ZTL hinges upon two fundamental light-regulated events. First, blue-light induces a conformational change within the N-terminal LOV domain that enhances LOV-mediated protein-protein interactions with GI^*26, 39*^. The resulting complex mutually stabilizes both proteins resulting in accumulation of ZTL and GI during the day. In conjunction with ZTL-GI complex formation, light inhibits the E3-ligase activity of ZTL, leading to decreased rates of degradation of ZTL targets PRR5 and TOC1^*9, 24, 25, 40-42*^. Throughout the night, decay of the C4a adduct activates ZTL mediated degradation activity allowing turnover of PRR5 and TOC1 in a time of day dependent manner^*9*^. How ZTL structural rearrangements, particularly in relation to the role of LOV-mediated dimers, impact degradation activity is currently unknown. Here, we observe that introduction of G46S resulted in two consequences 1) Stabilization of N- and C-terminal elements consistent with increased stability of ZTL in the light-state, and 2) The ZTL LOV domain undergoes a 180 degree rotation about the LOV dimer interface to favor a parallel orientation. We propose that rearrangement of the LOV dimer interface leads to a decrease in E3-ligase activity for light-state ZTL.

## Author Contributions

Protein expression, purification, and structural characterization of ZTL G46S and G46A constructs were conducted by AP, RG and BDZ. AB and NK conducted SAXS data analysis. Protein interaction studies in plant tissues were conducted by YHS and TI. All contributed to the writing of the manuscript.

## Acknowledgments

Work was funded by the National Institutes of Health (2R15GM109282 to BDZ, and R01GM079712 to TI) and by the Nuclear R&D Program of the Ministry of Science and ICT (MSIT), Republic of Korea (YHS). YHS and TI are also supported by Next-Generation BioGreen 21 Program (SSAC, PJ013386, Rural Development Administration, Republic of Korea).

---

**Table 1.** Data collection and refinement statistics (molecular replacement)

|  | G46S:G80R | G46A:G80R |
| --- | --- | --- |
|  | G46S:G80R | G46A:G80R |
| **Data collection** |  |  |
| Space group | I2(1)3 | I2(1)3 |
| Cell dimensions |  |  |
| *a*, *b*, *c* (Å) | 265.1, 265.1, 265.1 | 265.8, 265.8, 265.8 |
| $\alpha$, $\beta$, $\gamma$ (°) | 90, 90, 90 | 90, 90, 90 |
| Resolution (Å) * | 3.0 (3.11-3.00) | 3.0 (3.11-3.00) |
| *R*<sub>sym</sub> or *R*<sub>merge</sub> | 12.8 | 17.2 |
| *I* / σ*I* | 20.9 (2.9) | 17.5 (2.6) |
| Completeness (%) | 99.9 (99.9) | 99.7 (99.9) |
| Redundancy | 5.4 | 6.4 |
| CC1/2 | 0.987 (0.699) | 0.994 (0.642) |
| **Refinement** |  |  |
| Resolution (Å) | 3.0 | 3.0 |
| No. reflections | 61740 | 62237 |
| *R*<sub>work</sub> / *R*<sub>free</sub> | 19.3/22.2 (31.8/36.9) | 20.3/24.2 (31.0/35.2) |
| No. atoms |  |  |
| Protein | 7257 | 7250 |
| Ligand/ion | 237 | 229 |
| Water | 87 | 87 |
| *B*-factors |  |  |
| Protein | 56.5 | 52.1 |
| Ligand/ion | 50.9 | 45.5 |
| Water | 42.1 | 44.6 |
| R.m.s. deviations |  |  |
| Bond lengths (Å) | 0.016 | 0.016 |
| Bond angles (°) | 1.60 | 1.60 |

\*Highest-resolution shell is shown in parentheses.

