## Supplemental Figures for "Steric Interactions at Gln154 in ZEITLUPE Induce Reorganization of the LOV Domain Dimer Interface"

Supporting information:

Supporting Figure S1: Alternative assemblies in the crystal lattice and Gln154 orientations

Supporting Figure S2: SEC-SAXS data and analysis

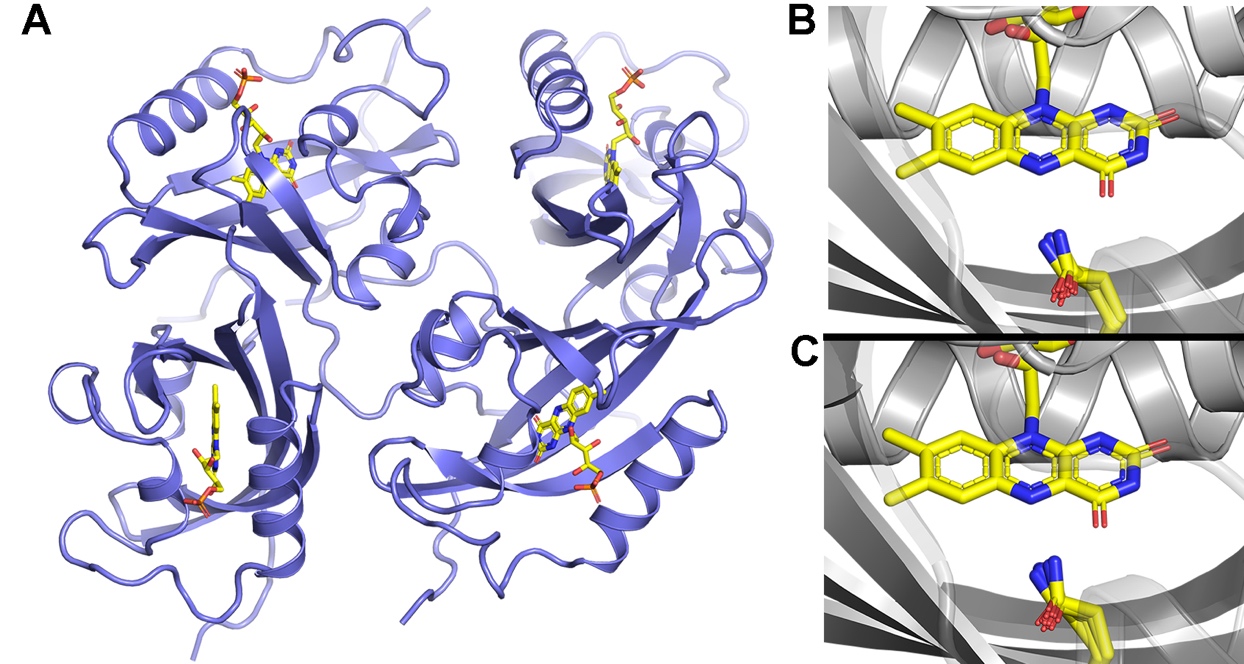

**Fig. S1:** G46S and G46A Structural Configurations. A) A tetrameric assembly present in the crystal structures. Dimeric, tetrameric, and the heptameric rings can be found within different selections of the asymmetric unit. Only the dimeric assembly has been observed in solution studies. B, C) All seven Gln154 orientations within the G46S (B) and G46A (C) crystal lattice. In all molecules the Gln side chain adopts a configuration that is partially buried to allow H-bonding interactions at N5 and O4 of the flavin isoalloxazine ring.

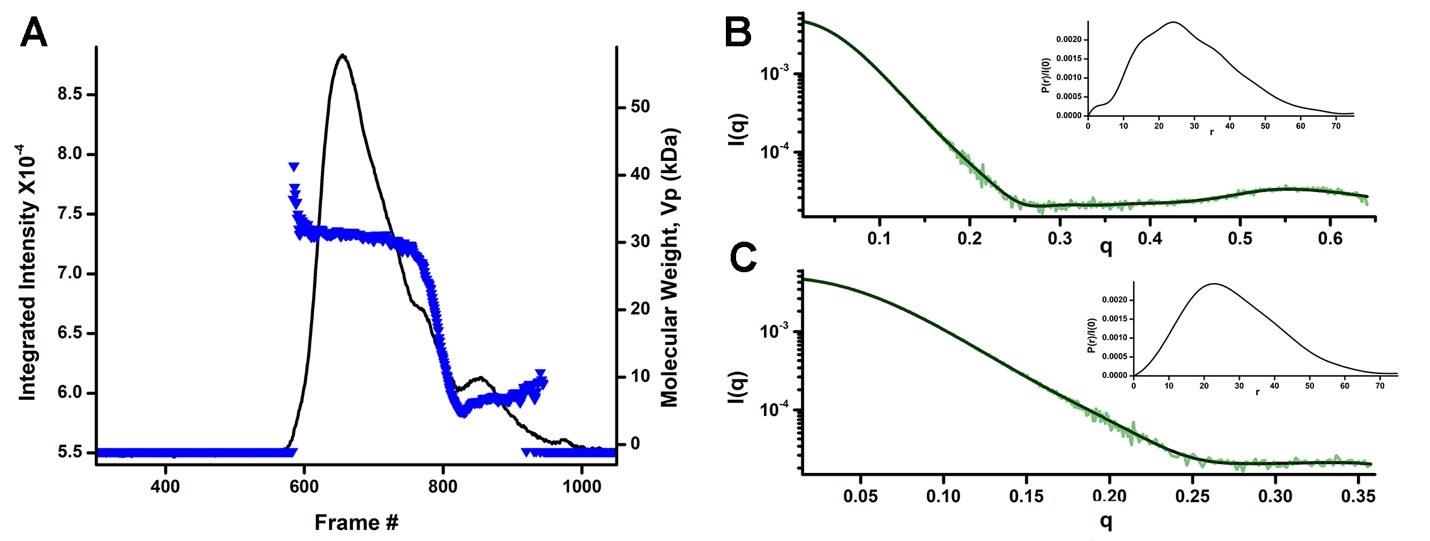

**Fig. S2:** SEC-SAXS of G46S:G80R ZTL. A) G46S:G80R elutes predominantly as a dimer with a secondary peak due to sample degradation. SAXS profiles were generated using frames 631-696 to capture the primary dimer peak. B) Full scattering profile for G46S:G80R from merged SAXS-WAXS data (olive) and corresponding fit of the pairwise distribution function (black line). Pairwise distribution profiles (inset) were generated using GNOM. C) Scattering profile truncated to q_max=8/Rg for DAMMIN reconstructions (olive) and corresponding fit of the pairwise distribution function (black line). Pairwise distribution profiles (inset) were generated using GNOM.
